# mapquik: Efficient low-divergence mapping of long reads in minimizer space

**DOI:** 10.1101/2022.12.23.521809

**Authors:** Barış Ekim, Kristoffer Sahlin, Paul Medvedev, Bonnie Berger, Rayan Chikhi

## Abstract

DNA sequencing data continues to progress towards longer reads with increasingly lower sequencing error rates. We focus on the critical problem of mapping, or aligning, low-divergence sequences from long reads (PacBio HiFi) to a reference genome, which poses challenges in terms of accuracy and computational resources when using cutting-edge read mapping approaches that are designed for all types of alignments. A natural idea would be to optimize efficiency with longer seeds to reduce the probability of extraneous matches; however, contiguous exact seeds quickly reach a sensitivity limit. We introduce `mapquik`, a novel strategy that creates accurate longer seeds by anchoring alignments through matches of *k* consecutively-sampled minimizers (*k*-min-mers) and only indexing *k*-min-mers that occur once in the reference genome, thereby unlocking ultra-fast mapping while retaining high sensitivity. We demonstrate that `mapquik` significantly accelerates the seeding and chaining steps — fundamental bottlenecks to read mapping — for both the human and maize genomes with *>* 96% sensitivity and near-perfect specificity. On the human genome, `mapquik` achieves a 30× speed-up over the state-of-the-art tool `minimap2`, and on the maize genome, a 350× speed-up over `minimap2`, making `mapquik` the fastest mapper to date. These accelerations are enabled not only by minimizer-space seeding but also a novel heuristic 𝒪(*n*) pseudo-chaining algorithm, which improves over the long-standing 𝒪(*n* log *n*) bound. Minimizer-space computation builds the foundation for achieving real-time analysis of long-read sequencing data.

## Introduction

Recent advances in DNA sequencing enable the rapid production of long reads with low error rates, e.g., Pacific Biosciences (PacBio) HiFi reads are 10 to 25 Kbp in length with ≤ 1% error rate. High-quality long reads have been used to accurately assemble genomes [20, 32, 9, 4]; complete the human genome [31]; accurately detect small variants in challenging genomic regions [33]; and further elucidate the landscape of large structural variants in human genomes [6]. Critical to these successes are algorithms that perform genomic data analysis, such as reconstructing a reference from reads (genome assembly) [25], or mapping reads to a reference genome (read mapping) [2]. With up to hundreds of gigabytes of sequenced data per sample, analysis algorithms need to balance efficiency with high sensitivity and accuracy (the percentage of reads mapped correctly), which is especially critical in rapid sequencing-to-diagnostics [35, 11].

We recently introduced the concept of minimizer-space computation [9], where only a small fraction of the sequenced bases are retained as a latent representation of the sequencing data, enabling orders of magnitude improvements in efficiency without loss of accuracy. Minimizers are sequences that are selected under some local or global minimality criteria [37, 41], similar to Locally Consistent Parsing (LCP), introduced by Şahinalp and Vishkin [5]. We applied the minimizer-space concept to perform genome assembly of long and accurate reads in minutes, instead of hours, and hypothesized that other types of genome analysis tasks would benefit from it in the future [9]. We now pursue the intuition that read mapping would also be amenable to minimizer-space computation, but there are multiple algorithmic challenges to overcome due to the repetitive nature of genomes, biological variation between samples and references, and sizable input data.

Two cornerstones of read alignment/mapping algorithms — ubiquitous in sequence analysis pipelines — are the *seeding* and *chaining* steps, where each read is locally placed at a homologous location in a reference genome. Seeding is carried out by finding pairs of matching seeds, which are snippets of DNA with high-confidence (exact or inexact) matches between a query and a reference genome. Seeds are initial matches that serve as anchoring points of alignments: They allow a challenging instance to be split into a set of easier sub-instances by aligning only the shorter intervals in-between seeds. In the short-read era, state-of-the-art alignment algorithms (e.g., `BWA-MEM` [22], `Bowtie2` [21], and `CORA` [44, 42]) typically relied on finding all possible seeds using a full-text index of the reference genome. For long reads, there has been a recent breakthrough by sampling and indexing only a relatively small number of short potential seeds from the reference genome, which has led to faster and more accurate mapping tools, e.g., `minimap2` [24] and `Winnowmap2` [16]. Chaining consists of finding maximal subsets of seeds that all agree on a certain genomic location [14]; seeds often have spurious matches due to their short lengths.

However, even the most recent long-read alignment tools are the bottlenecks in analysis pipelines. For instance, the popular `minimap2` software requires 12 CPU hours to map a typical PacBio HiFi dataset to the human genome, and `Winnowmap2` requires 15 CPU days, preventing both real-time analysis of sequencing data [26] and efficient reanalysis of previously-sequenced data collections [8]. A significant part of the `minimap2` and `Winnowmap2` running times are in their seeding and chaining steps [18]. These state-of-the-art long-read alignment tools are sensitive and accurate, but their underlying seed constructs (*k*-mers) are tailored to noisy reads. These small seed sizes induce longer computation times due to the multiple potential mapping locations of seeds that need to be examined and filtered out. Recent advances in short-read alignment methods have demonstrated that 98% of many organisms’ genomes are non-repetitive, and can be uniquely aligned to with longer seeds [7]. Therefore, it seems natural to explore the use of longer seeds also in long reads: this idea is at the heart of our approach.

Here, we provide a highly efficient read mapping tool for state-of-the-art and low-error long-read data. We introduce `mapquik`, which instead of using a single minimizer as a seed to a genome (e.g. `minimap2`), builds accurate longer seeds by anchoring alignments through matches of *k* consecutively-sampled minimizers (*k*-min-mers). Our approach borrows from natural language processing, where the tokens of the *k*-mers are the minimizers instead of base-pair letters. We demonstrate its 30× and 420× relative speedup over state-of-the-art mapping methods `minimap2` and `Winnowmap2`, respectively. We further apply `mapquik` to mapping simulated HiFi reads from the highly-repetitive maize genome, and demonstrate a 350× speed-up over `minimap2`, with,remarkably, no loss of sensitivity or accuracy.

One of our key conceptual advances is that minimizer-space seeds can serve as anchors in low-divergence alignments (e.g., from PacBio HiFi reads) via matches of *k* consecutively-sampled minimizers (*k*-min-mers), which results in orders of magnitude faster computation. This novel application of minimizer-space computation is entirely distinct from genome assembly, as no de Bruijn graph is constructed; this work establishes the versatility of *k*-min-mers in algorithms for biological sequences. We also show for the first time that indexing only the long minimizer-space seeds (*k*-min-mers) that occur uniquely in the genome is sufficient for sensitive and specific mapping. Another major conceptual advance is that by leveraging the high specificity of these seeds, we can devise a provably 𝒪(*n*) time (heuristic) pseudo-chaining algorithm, which improves upon the subsequent best 𝒪(*n* log *n*) runtime of all other known colinear chaining methods [14], without loss of performance in practice. We further study why simply using longer *k*-mers would be suboptimal. We believe that long seeds such as *k*-min-mers hold a promising future in ultra-fast long-read mapping, and beyond.

### Related work

`minimap2` [24] is a *de facto* standard for mapping accurate long reads to a reference genome. It applies a seed-and-extend strategy. Specifically, seeds are short *k*-mer minimizers, i.e., sequences of length *k* that are lexicographically-minimal within a window of *w* consecutive *k*-mers. The extension step is performed using an optimized implementation of the Needleman-Wunsch algorithm [30] with an affine gap penalty. Several attempts have been made to improve mapping performance compared to `minimap2`. `MashMap` [13] and `MashMap2` [15] compute read-versus-genome and genome-versus-genome mappings without an alignment step, and use 5× less memory than `minimap2` at the expense of longer runtime. In recent work developed concurrently to ours, unbeknownst to us, and not yet peer-reviewed, `BLEND` [10], a very recent aligner, uses strobemers [39] and locality-sensitive hashing (inspired by [5]) to speed up `minimap2` end-to-end by about 2×; however, their seeding approach is integrated into `minimap2`’s codebase, which is implemented and optimized for exact short seeds (minimizers [37]), thus suffering from similar limitations (for the sake of completeness, we compare to `BLEND` in Results).

Other works have focused on improving the sensitivity and accuracy of `minimap2`, at the expense of speed. `Winnowmap` [17] and `Winnowmap2` [16] use weighted minimizer sampling and minimal confidently-alignable substrings to better align in highly-repetitive regions, e.g., centromeres of chromosomes. `Winnowmap2` is around 15× slower than `minimap2` end-to-end, yet uses around 3× less memory.

In recent years, many research groups focusing on low-level and/or hardware-specific acceleration have proposed ways to accelerate `minimap2`. `mm2-fast` [18], a CPU acceleration of `minimap2` developed by Intel, achieved a 1.5× acceleration over `minimap2`, end-to-end on HiFi reads. The bulk of the speed-up was obtained in the alignment phase, not in the seeding/chaining phase (Figure 1 in [18]). The domain-specific language Seq was developed to speed up genomic sequence analysis, and achieved two orders of magnitude improvement in the homology table reconstruction of the `CORA` read mapper [44, 42].

**Fig. 1:**
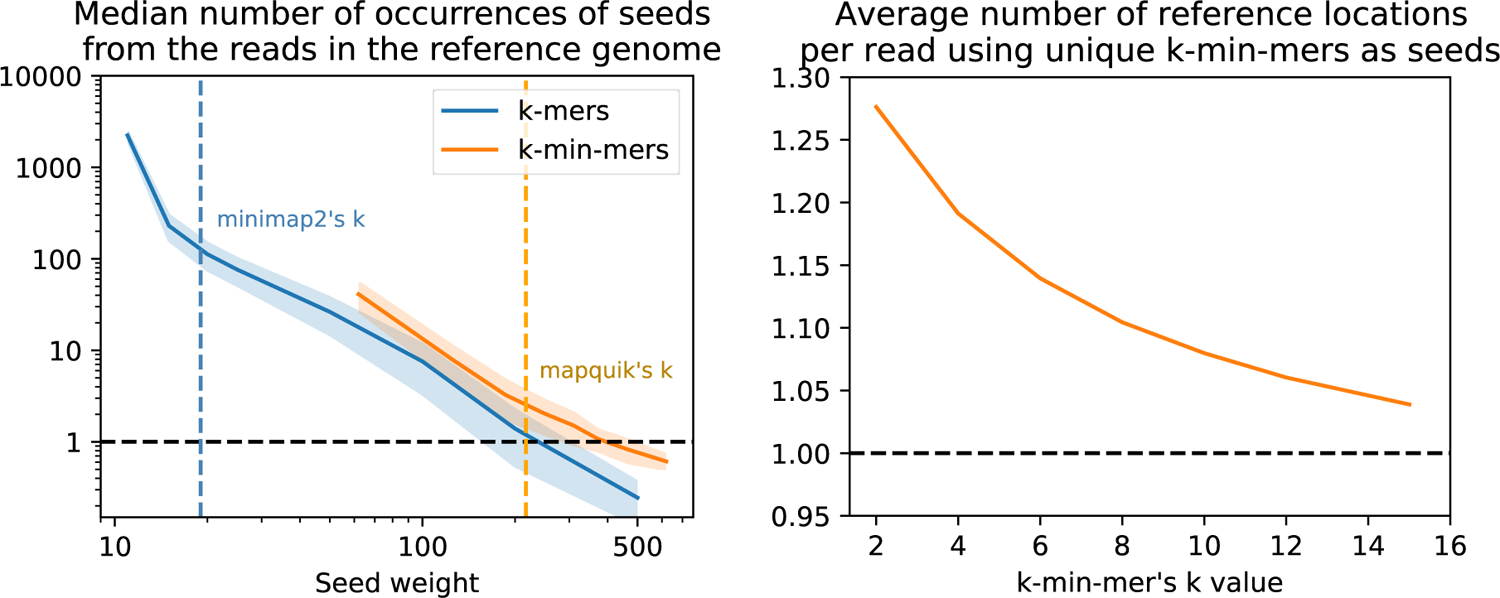
Increased sensitivity and specificity of *k*-min-mers versus long *k*-mers. Both panels use the human reference genome CHM13v2.0 and the HG002 DeepConsensus HiFi reads. Left panel: Each continuous line indicates the median abundance of read *k*-mers (shown in a darker blue line) and *k*-min-mers (shown in lighter orange line) in the reference, averaged across all reads (the closer to 1, the better). The vertical dashed darker blue line (respectively, the lighter orange line) corresponds to the seed length chosen by `minimap2` (respectively, by `mapquik`). The median is computed from a random sub-sample of 50,000 HG002 reads. Right panel: Average number of reference genome locations indicated by seed matches for each read using *k*-min-mers (the closer to 1 the better). *K*-min-mer parameters are *\** = 31, *δ* = 0.01 with *k* = 2 to 10 (left) and 2 to 20 (middle). Regular *k*-mer lengths are *k* = 12 to 500.

**Fig. 2:**
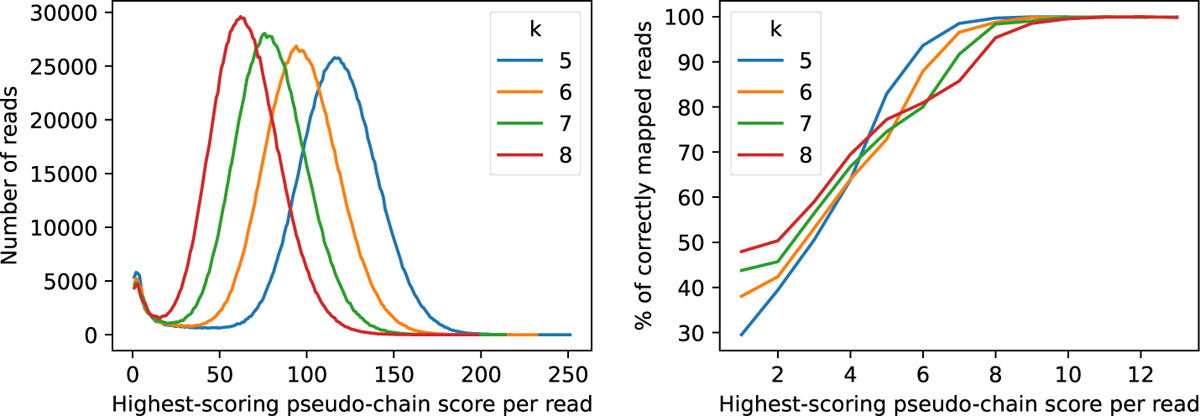
Effect of pseudo-chain score on mapping accuracy. The *x*-axis in both subfigures corresponds to the score of the maximal pseudo-chain per read; on the left, the *y*-axis denotes the total number of reads with the corresponding maximal pseudo-chain score, and on the right, the percentage of reads that are correctly mapped (assessed by `paftools_mapeval`) with the corresponding maximal pseudo-chain score. Only the scores below a threshold *s*, where *s* denotes the maximum score at which a read was mapped incorrectly, are plotted. The parameters used for `mapquik` were *k* = 5 to 8, *ℓ* = 31, *δ* = 0.01.

There also exist efficient implementations that employ specialized hardware. `mm2-ax` [38], a recent GPU acceleration of `minimap2` developed by NVIDIA, achieved a 2.5× – 5.4× acceleration over `mm2-fast` in the chaining step. Guo *et al*. also proposed GPU and FPGA accelerations of `minimap2` that respectively achieve 7× and 28× speed-ups on the distinct task of detecting pairwise read overlaps [12]. These specialized hardware accelerators are outside the scope of this work. Methods that accelerate short-read alignment, e.g., using cloud computing resources [40], or optimized *k*-mer indexing [1] are also not considered here, given that they do not support long reads.

### Why use *k*-min-mers as alignment seeds instead of *k*-mers?

Here, we motivate why *k*-min-mers are superior alignment seeds compared to *k*-mers for accurate long reads. Specifically, we formulate and verify the following two hypotheses: (1) Long exact *k*-mer seeds are inadequate for accurate long-read alignment due to lack of sensitivity, whereas (2) *k*-min-mers are adequate and also offer near-perfect specificity. An empirical analysis on simulated reads over the entire human genome (Figure 1) justifies these observations. The following experiments were performed using `Jellyfish` [27], `DSK` [36], `rust-mdbg` [9] and `mapquik`. All code and data are available in the `mapquik` repository (`experiments/figure-seeds/` folder).

The left panel of Figure 1 examines the specificity of *k*-mers and *k*-min-mers as seeds, by recording their number of occurrences in the CHM13v2.0 as a proxy for the number of potential mapping locations. The *x*-axis reports the *seed weight*, which for *k*-mers corresponds to their length, and for *k*-min-mers corresponds to *ℓ* × *k*, i.e., the total number of bases in the *k*-min-mer minimizers. As indicated by the plot, *k*-mer seeds either have too many matches in the reference (from tens to thousands in the *k* = 10 to 100 range), or too few (below one match for *k >* 300). Notably, `minimap2` uses a default *k* value of 19 for HiFi reads, reflecting that it has to sift through hundreds of false matches for each read on average. On the other hand, *k*-min-mers have orders of magnitude fewer potential matches to examine, on average one to tens depending on *k*, owing to their longer lengths being less affected by genomic repetitions.

The right panel of Figure 1 illustrates that selecting all the *k*-min-mers that are seen only once in the reference genome is a viable indexing strategy. Indeed, the reads have, on average, 1.05 – 1.30 candidate reference genome locations when all their *k*-min-mers are queried on such an index. This hints that a read mapping algorithm based on *k*-min-mers is likely to immediately find the right genome location by querying all read *k*-min-mers, and only paying attention to those that occur once in the genome. This algorithm potentially would not even require a subsequent chaining step, given the low number of false matches to remove. This is in stark contrast with existing *k*-mer-based algorithms, for which colinear chaining removes hundreds of false seed matches per read.

## Methods

### Minimizer-space read mapping with mapquik

We developed `mapquik`, a read mapper based on *k*-min-mer seeds, which allow computation in minimizer-space (Figure 3). `mapquik` follows a seed-and-extend strategy used by most read mappers, with two exceptions: (1) Only the *k*-min-mers that appear exactly once in all target sequences are indexed, and (2) unlike a typical colinear chaining procedure that makes use of a dynamic programming formulation, e.g., in `minimap2` [24], a linear-time recursive extension step is performed for each initial *k*-min-mer match between the query and the reference, followed by a novel, provably linear-time step we call *pseudo-chaining*, which ensures that *k*-min-mer matches are approximately colinear.

**Fig. 3:**
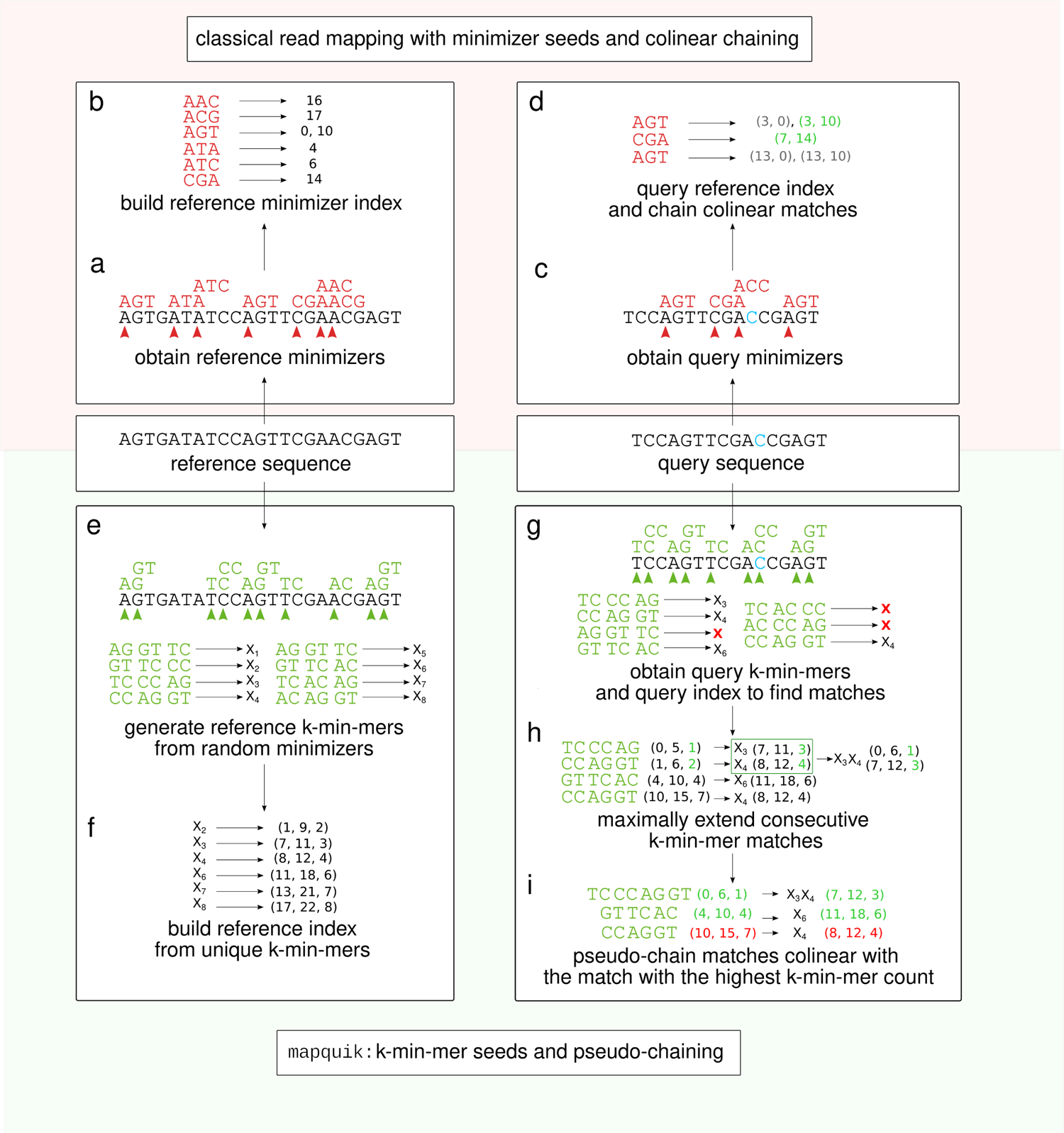
Overview of the long-read mapping pipeline using mapquik and comparison with state-of-the-art methods using minimizers as seeds. State-of-the-art read mappers such as `minimap2` and `Winnowmap2` (top) build an index for a reference sequence by computing window minimizers (*k* = 3, *w* = 5) (a), and storing the positions of the minimizers in the index (b). In order to map a query sequence using the reference index (top right, nucleotide `C` in blue denotes a sequencing error), mappers compute the minimizers on the query sequence (c), and find matches between the minimizers of the query and those in the reference index. Once minimizer matches are found, `minimap2` and `Winnowmap2` perform a colinear chaining step to output a high-scoring set of matches, using dynamic programming (d). In contrast, `mapquik` (bottom) indexes reference sequences by generating *k*-min-mers, *k* consecutive, randomly-selected minimizers of length *ℓ* (*k* = 3, *ℓ* = 2) (e), and storing only the *k*-min-mers that appear exactly once in the reference (f). `mapquik` stores the start and end position of each *k*-min-mer, along with the order the *k*-min-mers appear in. In order to map a query sequence using the *k*-min-mer index, `mapquik` first obtains matches between the query and the reference index by querying the index with each query *k*-min-mer (g). *k*-min-mer matches are extended if the next immediate pair of *k*-min-mers also match (h). Instead of a colinear chaining step, `mapquik` performs a linear-time pseudo-chaining step, which locates matches that are colinear with the match with the highest number of *k*-min-mers (i).

The philosophy behind these drastic changes is that mapping long and accurate reads to close reference genomes is “easy enough” that long minimizer-space seeds will be sufficient for the vast majority of the reads. The remaining few unmapped reads may be fed to a more sensitive, albeit slower, read mapper such as `minimap2` or `Winnowmap2` (see Discussion).

### Algorithmic details

Figure 3 and Algorithm 1 describe the steps of `mapquik`. The following subsections go into more detail for each step.

#### Algorithm 1 The mapquik algorithm

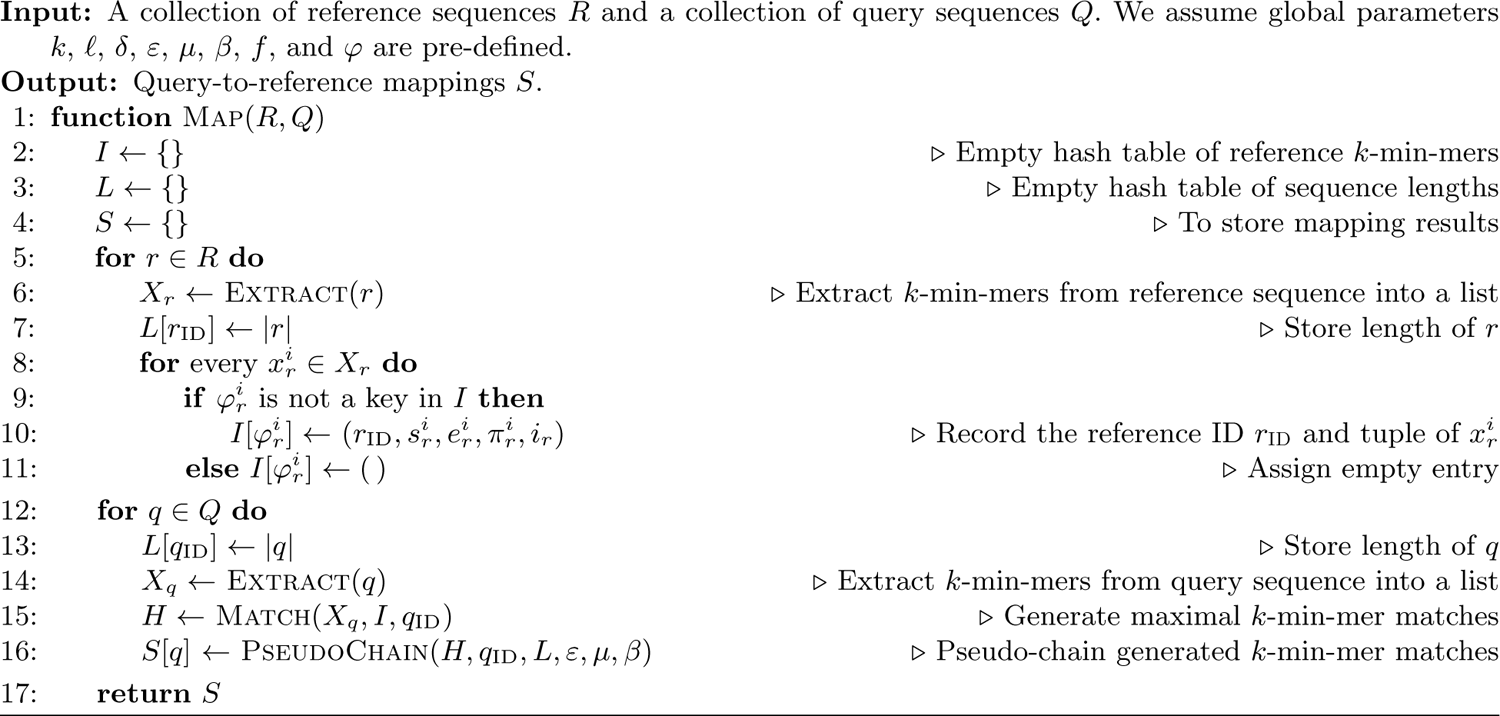

### Methodological formalization

For a fixed integer *ℓ >* 1, let *f*: *Σ*^*ℓ*^ ⟼ [0, *H*] be a random hash function that maps strings of length *ℓ* to integers between 0 and *H*. In practice, we use a 64-bit hash function. Moreover, we require *f* to be invariant with respect to reverse-complements, i.e., an *ℓ*-mer and its reverse complement map to the same integer. For a density 0 < *δ* < 1, we define *U*_*ℓ,δ*_ as the set of all *ℓ*-mers *m* with *f* (*m*) < *δ* · *H*. We refer to the elements of *U*_*ℓ,δ*_ as (*ℓ, δ*)-*minimizers*. Note that whether an *ℓ*-mer is a (*ℓ, δ*)-minimizer does not depend on any sequence besides the *ℓ*-mer itself.

Let *S* be a sequence of length ≥ *ℓ*. We define its *minimizer-space representation M*_*S*_ as an ordered list of (*ℓ, δ*)-minimizers that appear in *S*. Since the contents of *U*_*ℓ,δ*_ only depend on *f*, and as long as the same hash function *f* is used, *M*_*S*_ will always be a subset of *U*_*ℓ,δ*_. We omit *S* from the subscript when it is obvious from the context.

Let *M* be a minimizer-space representation of *S* and let *k >* 0 be a fixed integer parameter. We define a *k-min-mer x*^*i*^ of *S* as an ordered list of *k* consecutive minimizers in *M* starting from index *i*, i.e., *x*^*i*^ = (*m*_*i*_, …, *m*_*i*+*k*−1_). We denote the ordered list of all *k*-min-mers (*x*^0^, …, *x*^|*M*|−*k*^) of *S* as *X*_*S*_. We omit *S* from the subscript when it is obvious from the context.

In order to avoid explicitly storing nucleotide sequences of minimizers, we use a random hash function *φ* that maps sequences of *k ℓ*-mers to 64-bit hash values. We define *φ* so that it is invariant to reversing the order of the *k*-min-mer, i.e., *φ*(*x*^*i*^) = *φ*(rev(*x*^*i*^)), where rev(*x*^*i*^) denotes the list of minimizers of *x*^*i*^ with the order reversed. This is achieved by hashing *x*^*i*^ and rev(*x*^*i*^), and taking *φ*(*x*^*i*^) to be the minimum value.

Then, instead of storing *X* as an ordered list of *k*-min-mer sequences, we store the *i*^th^ *k*-min-mer *x*^*i*^ of *X* as a tuple (*i, φ*^*i*^, *s*^*i*^, *e*^*i*^, *π*^*i*^), where

– *i* is the rank of *x*^*i*^ in *X*,
– *φ*^*i*^ = *φ*(*x*^*i*^),
– *s*^*i*^ and *e*^*i*^ the nucleotide start and end positions of *x*^*i*^, i.e., the start position of minimizer *m*_*i*_ and the end position of minimizer *m*_*i*+*k*−1_, respectively, on *S*,
– *π*^*i*^ a Boolean variable that evaluates to 1 if, in the construction of *φ*, the hash of *rev*(*x*^*i*^) was smaller than that of *x*^*i*^, and 0 otherwise.

We call this the *tuple of x*^*i*^. When the sequence *S* is not obvious from the context, we add it as a subscript in the above notation, e.g., 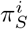. The hash functions *f* and *φ* are instantiated prior to constructing the *k*-min-mer lists for the input sequences. We consistently use the same functions *f* and *φ* when selecting minimizers and consequently building *k*-min-mer lists for both the reference and query sequences throughout.

### Indexing reference sequences

The `mapquik` index *I* is a hash table that associates a *k*-min-mer *x* to its unique position in the reference genome, whether or not it appears reverse-complemented, and its rank in the list of *k*-min-mers *X*. As described in Section 1, this information is represented by a tuple in the form (*r*_ID_, *s*^*i*^, *e*^*i*^, *π*^*i*^, *i*). We construct *I* using a two-pass approach. First, we call the Extract function. It builds the list *X* by a linear scan through the reference sequence, during which it identifies minimizers and outputs each *k* consecutive minimizers, together with their hash values. We use the same efficient algorithm as in [9] (which runs in 𝒪(|*X*|) time), so we do not include the pseudo-code for Extract here. Second, we load every entry of *X* into a hash table *I*, indexed by the hash value. In this step, we discard from *I* any *k*-min-mer that appears in more than one reference location. Therefore, the hash table holds a single value per distinct *k*-min-mer key.

### Locating and extending query-to-reference *k*-min-mer matches

Informally, a *k-min-mer match* is a stretch of *k*-min-mer seeds which appear consecutively both in the reference and the query (under the hash function used). Note that all matches are unique, in the sense that all seed *k*-min-mers appear only once in the genome, by definition. Given a query *q* and a reference *r*, we formally define a *match* as a triple (*i, j, c*) such that 0 ≤ *i* ≤ |*X*_*q*_| − *c*, 0 ≤ *j* ≤ |*X*_*r*_| − *c, c* ≥ 1, and, for all 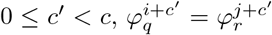. A match (*i, j, c*) is said to be *maximal* if cannot be further extended to the right or left, i.e., (1) either *i* = 0, *j* = 0, or 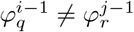, and (2) either *i* + *c* = |*X*_*q*_|, *j* + *c* = |*X*_*r*_|, or 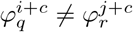.

For each query *q*, the `mapquik` algorithm first builds the list of *k*-min-mers *X*_*q*_, sorted in increasing order of location (line 14). Then, it runs the Match routine (line 15), which finds all maximal matches between *q* and the reference. Match works by scanning through *X*_*q*_ and, for each seed *x* ∈ *X*_*q*_, using the reference index *I* to see if *x* exists in the reference. If it does, then it marks the start of a match and proceeds to extend the match to the right as long as the seeds continue to match. Since we only have to query the index with the hash value of each *k*-min-mer in *X*_*q*_, the extension procedure can be done during a single linear pass over the elements of *X*_*q*_, and thus takes 𝒪(|*X*_*q*_|) time (assuming 𝒪(1) hashing, look-ups, and insertions). Care must be taken due to reverse complements, which can change the direction of matching, but we omit these details. For completeness, the full algorithm is in the Appendix (Algorithm 2).

In theory, generating a single 64-bit hash value for each unique *k*-min-mer could lead to hash collisions and, consequently, lead to false *k*-min-mer matches. However, since a *k*-min-mer match can only be extended with a consecutive *k*-min-mer match, *k*-min-mers that match an entry in the reference due to a hash collision are likely to be singletons and get filtered out in the pseudo-chaining step, with no decrease in final accuracy.

### From maximal *k*-min-mer matches to pseudo-chains

Recall that *k*-min-mer matches are extended based solely on whether the next immediate *k*-min-mer of *q* matches the next immediate *k*-min-mer of *r*. However, *k*-min-mer matches on *q* might occur in multiple non-overlapping positions on *r* (due to sequencing errors or biological variation in *q*), i.e., for two matches (*i, j, c*) and (*i*′, *j*′, *c*′) between *q* and *r*, it is not necessarily true that |*i* − *j*| = |*i*′ − *j*′|.

In order to output a list of matches that are likely to be true positives while avoiding a computationally-expensive dynamic programming procedure, `mapquik` uses a *pseudo-chaining* procedure which finds *k*-min-mer matches between a query *q* and a reference *r* that are gap-bounded colinear, but not all pairwise colinear.

Concretely, let *h* = (*i, j, c*) and *h*′ = (*i*′, *j*′, *c*′) be two matches, and consider the coordinates (*s*_*q*_, *e*_*q*_, *π*) and (*s*_*r*_, *e*_*r*_, *π*) of respectively the first and last *k*-min-mers of *h*, and 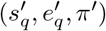 and 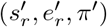 for the *k*-min-mers of *h*′. Let *g >* 0 be a fixed-integer gap upper bound. We say that *h* and *h*′ are *gap-bounded colinear* if

– the matches are on the same relative strand, i.e., *π* = *π*′,
– the reference start positions of the matches agree with the order of the matches, i.e., if 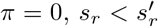; or if 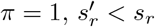, and
– the length of the gap between the two matches in the query is similar to that on the reference, i.e., if 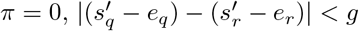; or if 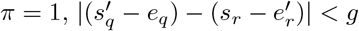.

This last *gap length difference* condition, on top of the traditional definition of colinearity (the first two conditions), ensures that the regions outside the matches are similar in length. A similar parameter (*ϵ*) is used in `minimap` [23].

Let *H* be the list of all maximal *k*-min-mer matches between a read *q* and a reference sequence *r*. We define a *pseudo-chain Ψ*^*i*^ as the list of all matches in *H* that are colinear with the *i*^th^ match in *H*; we say that *Ψ*^*i*^ is *anchored at i*. Note that even though every match in *Ψ*^*i*^ is colinear with the *i*^th^ match in *H*, it is not necessarily true that every pair of matches in *Ψ*^*i*^ are pairwise colinear, thus *Ψ*^*i*^ does not satisfy the criteria of chains as defined in other works (e.g., [24]).

The *score* of a pseudo-chain *Ψ*^*i*^ is the number of matching *k*-min-mers in *Ψ*^*i*^, i.e.,

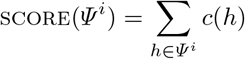

where *c*(*h*) denotes the number of matching *k*-min-mers in match *h*. Since the maximal matches in *Ψ*^*i*^ are guaranteed to not share any query *k*-min-mers because both the start and end locations of each maximal match are distinct, the cumulative sum of the number of matching *k*-min-mers in each match in *Ψ*^*i*^ equals the number of total matching *k*-min-mers in *Ψ*^*i*^.

### Computing high-scoring pseudo-chains in linear time

We now introduce a novel algorithm for computing a single high-scoring pseudo-chain *Ψ*\* for a query *q* given a list of maximal matches, and prove that it runs in 𝒪(*n*) time. The match extension step outputs a hash table *H* of maximal *k*-min-mer matches per reference, indexed by their reference identifier. In the pseudo-chaining step, however, the objective is to output a single list of matches between *q* and a single reference, even though *H* might contain matches between *q* and more than one reference sequence. We first initialize *Ψ*\* = [], and iterate over the key-value tuples in *H*, processing each list of maximal matches *H*_*q,r*_ for a single reference *r* one by one. In every iteration, we obtain a candidate pseudo-chain *Ψ*_*q,r*_ from the list of maximal matches *H*_*q,r*_ by computing the pseudo-chain anchored at the match in *H*_*q,r*_ with the highest number of matching *k*-min-mers. After computing *Ψ*_*q,r*_, we compare its score to that of *Ψ*\*, and replace *Ψ*\* with *Ψ*_*q,r*_ if score(*Ψ*_*q,r*_) *>* score(*Ψ*\*). At the end of the loop, *Ψ*\* will be the highest-scoring pseudo-chain out of all possible candidate pseudo-chains per reference sequence in *H*.

Finally, if the pseudo-chain *Ψ*\* has score ≥ *µ* or length ≥ *β*, where *µ* and *β* are user-defined parameters, we retrieve the query and reference coordinates of the region covered by the matches in *Ψ*\*. The final query and reference coordinates for a mapping between query *q* and reference *r* is computed by extending the start and end coordinates of the first and last matches in *Ψ*\* to the length of the query. In the Appendix, Algorithm 3 provides a complete description of the pseudo-chaining procedure, and Algorithm 4 describes the coordinate computation step.

#### Proof of pseudo-chaining algorithm’s complexity

The complexity of computing pseudo-chain *Ψ*^*i*^ for each read *q* is as follows. Let *n* be the total number of matches in *H*. To determine *Ψ*\*, each candidate pseudo-chain *Ψ*_*q,r*_ for a single reference sequence *r* needs to be computed. Computing a single pseudo-chain *Ψ*_*q,r*_ requires determining the match with the highest number of matching *k*-min-mers and comparing each match in *H*_*q,r*_ to this match, which can both be performed in *Θ*(|*H*_*q,r*_|) time. Moreover, every single candidate pseudo-chain (for every reference in *H*) needs to be computed to determine *Ψ*\*. Then, the running time of the pseudo-chaining procedure is *Θ*(∑_*r*∈*H*_ |*H*_*q,r*_|). Note that |*H*_*q,r*_| ≤ *n*, and the number of reference sequences that appear in *H* is upper bounded by the total number of reference sequences, which is 𝒪(1). Hence, the pseudo-chaining procedure runs in 𝒪 (*n*) time, where *n* is the total number of matches in *H*.

Note that colinear chaining (as implemented by state-of-the-art read mappers) has an asymptotic complexity of 𝒪 (*n* log *n*). We also implemented two alternative heuristics that (1) computes *c* pseudo-chains anchored at *c* matches with the highest number of *k*-min-mers (thus running in 𝒪(*cn*) time), and (2) sets *c* = *n* and computes all possible pseudo-chains (thus running in 𝒪(*n*^2^) time). However, we observed that the runtime of the 𝒪(*n*) pseudo-chaining procedure is faster in practice: In our tests, the 𝒪(*n*) pseudo-chaining procedure performed ∼ 20% − 50% faster than the other heuristics, with little decrease in accuracy.

## Results

### Datasets and mapping evaluation

We used the complete human reference genome CHM13v2.0 for our evaluations. We constructed a simulated dataset of long reads with 99% base-level accuracy and 24 Kbp mean length using `pbsim` [34], mimicking HiFi reads at 10× genome coverage. We also used real HiFi reads for the HG002 individual corrected using DeepConsensus [3], at 30× genome coverage. For maize, we simulated reads from the maize RefSeq genome (`GCF_902167145.1`) at 30× coverage using the same protocol as the human simulated reads. `mapquik` was run with default parameters (*k* = 5, *ℓ* = 31, *δ* = 0.01, *β* = 4, *µ* = 11, *ε* = 2000). All other tools were run with default parameters in HiFi mapping mode. Command lines and versions are given in the Appendix.

For simulated reads, we assessed mapping accuracy using the `mapeval` command of the `paftools` software distributed in the `minimap2` package. A read is considered to be correctly mapped if the intersection between the true and mapped reference intervals is at least 10% of their union. For real reads, we evaluated the concordance between the alignments of `minimap2` and `mapquik` using a custom script (`experiments/intersect_pafs.py` in the GitHub repository), similar to `mapeval`.

Mappers report a *mapping quality* metric for each read, indicating their confidence that the read is mapped at the right location, as an integer between 0 and 60 where 60 corresponds to the highest confidence. In our evaluation of both the simulated and real datasets, we focus on reads with the highest mapping quality (Q60), as reads with low mapping quality are less frequent and are often removed in downstream applications (e.g., the popular variant calling pipeline GATK [28] filters out reads with mapQ ≤ 20 by default). Furthermore on `minimap2` results, `mapeval` only reported two mapping quality groups (Q0 and Q60), with no value in-between. In `mapquik`, we report a mapQ of 60 if the output pseudo-chain has score ≥ *µ* or length ≥ *β*, and 0 otherwise.

### mapquik achieves faster and accurate mapping of HiFi reads to the human genome

Table 1 shows the overall performance of `mapquik` and other evaluated methods (`minimap2`, `mm2-fast`, `Winnowmap2`, and concurrently-developed `BLEND`) on mapping simulated and real PacBio HiFi reads to the human genome and the maize genome. On the simulated human dataset, `mapquik` is 32× faster than `minimap2`. `mapquik` maps 97.2% of reads at a mapQ score of 60 (Q60), indicating a high-confidence match, with 2 errors. In contrast, other mappers map 97.4 – 99.0% of reads at Q60 with also no/almost no error.

**Table 1:** **Mapping statistics** of `mapquik` and other evaluated methods (`minimap2`, `mm2-fast`, `Winnowmap2`, `BLEND`) on simulated and real HiFi human reads, and simulated maize HiFi reads. Only reads with reported mapping quality over 60 were included in columns 1 and 3. Incorrectly-aligned reads were detected using `paftools_mapeval`. “Time” column consists of wall-clock times and includes on-the-fly reference indexing. Reads were un`gzip`ped. Tools were run on 10 threads. For `Winnowmap2`, the time for reference *k*-mer counting (`meryl`) was not included. The last column indicates the speed-up over `minimap2` taken as a baseline.

| Tool name | Mapped Q60 | Q < 60 or unmapped | Wrong Q60 | Memory (GB) | Runtime (s) | Speed-up |
| --- | --- | --- | --- | --- | --- | --- |
| CHM13 10× coverage **simulated** 24 kbp HiFi reads |  |  |  |  |  |  |
| **minimap2** | 1,475,985 | 14,425 | 0 | 10.6 | 762 | 1x |
| **mm2-fast** | 1,475,985 | 14,425 | 0 | 10.6 | 709 | 1.07x |
| **Winnowmap2** | 1,474,449 | 15,961 | 5 | 12.2 | 10,516 | 0.07x |
| **BLEND** | 1,452,658 | 37,752 | 1 | 5.9 | 165 | 4.62x |
| **mapquik** | 1,448,711 | 41,699 | 2 | 12.1 | 24 | 31.75x |
| HG002 30× coverage **real** 24 kbp HiFi reads (DeepConsensus) |  |  |  |  |  |  |
| **minimap2** | 3,611,990 | 303,304 | N/A | 10.6 | 3,081 | 1x |
| **mm2-fast** | 3,611,983 | 303,311 | N/A | 10.6 | 2,635 | 1.17x |
| **Winnowmap2** | 3,835,225 | 80,069 | N/A | 8.8 | 83,180 | 0.04x |
| **BLEND** | 3,708,582 | 206,712 | N/A | 6.0 | 615 | 5.01x |
| **mapquik** | 3,760,677 | 154,617 | N/A | 12.1 | 105 | 29.34x |
| Maize 30× coverage **simulated** 24 kbp HiFi reads |  |  |  |  |  |  |
| **minimap2** | 2,807,058 | 58,730 | 1 | 5.7 | 16,801 | 1x |
| **mm2-fast** | 2,807,059 | 58,729 | 1 | 5.6 | 13,951 | 1.2x |
| **Winnowmap2** | 2,854,043 | 11,745 | 93 | 13.4 | 15,440 | 1.09x |
| **BLEND** | 2,836,244 | 29,544 | 5 | 4.3 | 333 | 50.45x |
| **mapquik** | 2,837,524 | 28,264 | 2 | 13.2 | 48 | 350.02x |

On the real dataset, a similar trend is observed, and `mapquik` outperforms all mappers except `Winnowmap2` in terms of percentage of reads mapped (96.1%), as opposed to 92.2 – 97.9% for the other tools. The concordance of `minimap2` and `mapquik` mappings is 99.8% on the Q60 reads mapped by `minimap2`. All mappers required less than 13 gigabytes of memory on the human genome (`MashMap2` was not further evaluated as it took over 13 wall-clock hours on the simulated human dataset, and does not output mapping quality scores).

A highlight of `mapquik` is a 350× mapping speed-up compared to `minimap2` on the maize genome. Remarkably, this speed-up comes with no loss of sensitivity, as `mapquik` reports the second highest number of mapped reads at mapping quality 60 across all tools after `Winnowmap2`, with near-perfect precision. `Winnowmap2` is faster than its performance on the human genome and that of `minimap2` on the maize genome.

We further investigated why some reads were mapped at lower qualities than Q60 or were not mapped at all. Out of 42,198 reads from the simulated human dataset that were not mapped at Q60 by `mapquik`, 94.4% of these reads intersected with centromeric/satellite regions of chromosomes, as reported by `bedtools` using the `chm13v2.0_censat_v2.0.bed` annotation from the T2T consortium. Thus, the vast majority of reads not aligned at Q60 corresponds to challenging genomic regions that would likely have been masked in downstream analyses, making their lack of alignment potentially inconsequential. We hypothesize that similar conclusions hold on real data, but this cannot be ascertained as the true reference interval of each read is not known.

### Efficient genome indexing using *k*-min-mers

The `mapquik` seeding strategy provides ultra-fast construction of an index that records unique *k*-min-mer positions across the reference genome; this index is of independent interest. Table 2 demonstrates the computing resources necessary to index a human genome, compared to `minimap2`, `mm2-fast`, `Winnowmap2` and the concurrently-developed `BLEND`.

**Table 2:** **Indexing a reference human genome** using `mapquik` and other evaluated methods (`minimap2`, `mm2-fast`, `Winnowmap2`, `BLEND`). The CHM13v2.0 reference sequence was given as input to each tool. Tools were run using 10 threads with warm cache, i.e., the reference genome file was already pre-loaded in memory. The “Indexed sequences” column indicates the number of distinct sequences that are keys of the final index. The “Singletons” column indicates how many indexed sequences have only one position in the reference genome.

| Tool name | Indexed sequences | Singletons | Memory (GB) | Wall-clock (seconds) |
| --- | --- | --- | --- | --- |
| **minimap2** | 215,125,355 | 92% | 10.1 | 33 |
| **mm2-fast** | 215,125,355 | 92% | 10.1 | 28 |
| **Winnowmap2** | 23,616,987 | 41% | **2.6** | 64 |
| **BLEND** | 111,799,540 | 97% | 5.3 | 23 |
| **mapquik** | 39,603,738 | 100% | 12.1 | **4.6** |

Indexes created by those mapping methods are different in nature. Yet, a similar order of magnitude of sequences (tens to hundreds of millions) end up being indexed. `minimap2` and `mm2-fast` index positions of windowed minimizers. Those minimizers are different from the (*ℓ, δ*)-minimizers defined in this article. `Winnowmap2` indexes weighted minimizers to increase the accuracy of seeds. `BLEND` (in HiFi mapping mode) indexes a locality-sensitive seed built from strobemers [39], which are chains of consecutive windowed minimizers [37], different in nature from *k*-min-mers.

`mapquik` indexing is 7× faster than `minimap2` and 5× faster than `BLEND`. We anticipate that this index will have other uses beyond long-read mapping, such as large-scale sequence search. We have already demonstrated the usefulness of performing *k*-min-mer search in anti-microbial resistance tracking [9], albeit in earlier work that did not benefit from the speed-up of such an index presented here.

### Comparison with BLEND

While concurrently-developed and not yet peer-reviewed, we compare to `BLEND` in Tables 1 and 2 for completeness, and demonstrate that `mapquik` achieves a 6× speed-up over it on human data.

### Limitations of our study

We evaluate the limitations of the method with respect to several aspects: The choice of *k*, internal cut-off parameters, and the requirement of having low divergence between the reads and the reference.

As seen in Table 1, `mapquik` is unable to map some of the reads, mostly in low-complexity regions of the genome, partially because no indexed *k*-min-mer exists, but also because of the lack of any long enough (i.e., high-scoring) pseudo-chain. Here, we examine the magnitude of the second effect. Figure 2 (left) shows the total number of reads (*y*-axis) whose highest pseudo-chain score is given on the *x*-axis. Read numbers follow approximately a Gaussian distribution, confirming the soundness of filtering out the leftmost tail containing erroneous pseudo-chains. Figure 2 (right) investigates filtering thresholds by showing the percentage of reads mapped correctly (*y*-axis) at each pseudo-chain score per read (*x*-axis). The monotonically-increasing relationship suggests that the pseudo-chain score is a reliable proxy for evaluating mapping accuracy. Mapping accuracy plateaus around pseudo-chain scores of 9 to 11. In our implementation, we apply a threshold of pseudo-chain score ≥ *µ* with *µ* = 11 by default (see Algorithm 3 in the Appendix).

Supplementary Figure 4 shows that the mapping performance of `mapquik` degrades markedly when iden-tity between reads and the reference is lower than 97%, and less than 1% of the reads are mapped at Q60 for identities below 93%. Therefore, `mapquik` is not suitable for mapping PacBio CLR reads, and potentially also Oxford Nanopore reads until base-calling consistently reaches identity levels above 98%. Remarkably, the mapping error rate at Q60 remains negligible at all identity levels.

## Discussion

We have demonstrated that long-read mapping can be sped up by over an order of magnitude by newly substituting short *k*-mer seeds with *k*-min-mers that occur uniquely in the genome. We have implemented this strategy in a novel read mapper, `mapquik`, and shown its superior performance as compared to other state-of-the-art tools. As sequencing reads are getting longer and more accurate, we anticipate that our approach will particularly further benefit from technological advances: Longer reads will be increasingly easier to map with minimizer-space seeds. Our approach demonstrates that minimizer-space computation can be successfully applied to read mapping and overcomes a significant barrier for real-time analysis of sequencing data.

A potential concern with providing a faster alignment method is the loss of sensitivity in hard-to-map regions, such as centromeres or structural variant breakpoints. One could partially mitigate this concern by performing a conservative, but fast alignment of reads using `mapquik`, and remapping the unmapped reads with `minimap2` or `Winnowmap2` to increase alignment sensitivity, while keeping the efficiency of `mapquik`.

Future extensions of this work include implementing base-level alignment, which will allow the design of a complete single-nucleotide variant calling pipeline, as well as structural variant calling built on top of `mapquik`. Since the *k*-min-mer matches have *k* exact matches of minimizers of length *ℓ*, only the regions in-between minimizers and in-between neighboring *k*-min-mer matches would need to be aligned, which potentially lowers both the memory usage and runtime of the alignment step. Another potential improvement to mapquik would be to refine the mapping quality scores, in light of the observations made in Figure 2.

## Software availability

https://github.com/ekimb/mapquik

## Acknowledgements

The authors thank Erik Garrison, Younhun Kim, Victoria Popic, and Rohit Singh for valuable feedback.

## Appendix

### A.1 Algorithms

Algorithms for specific steps of `mapquik` are given here for completeness.

#### Algorithm 2 Initiating and extending *k*-min-mer matches

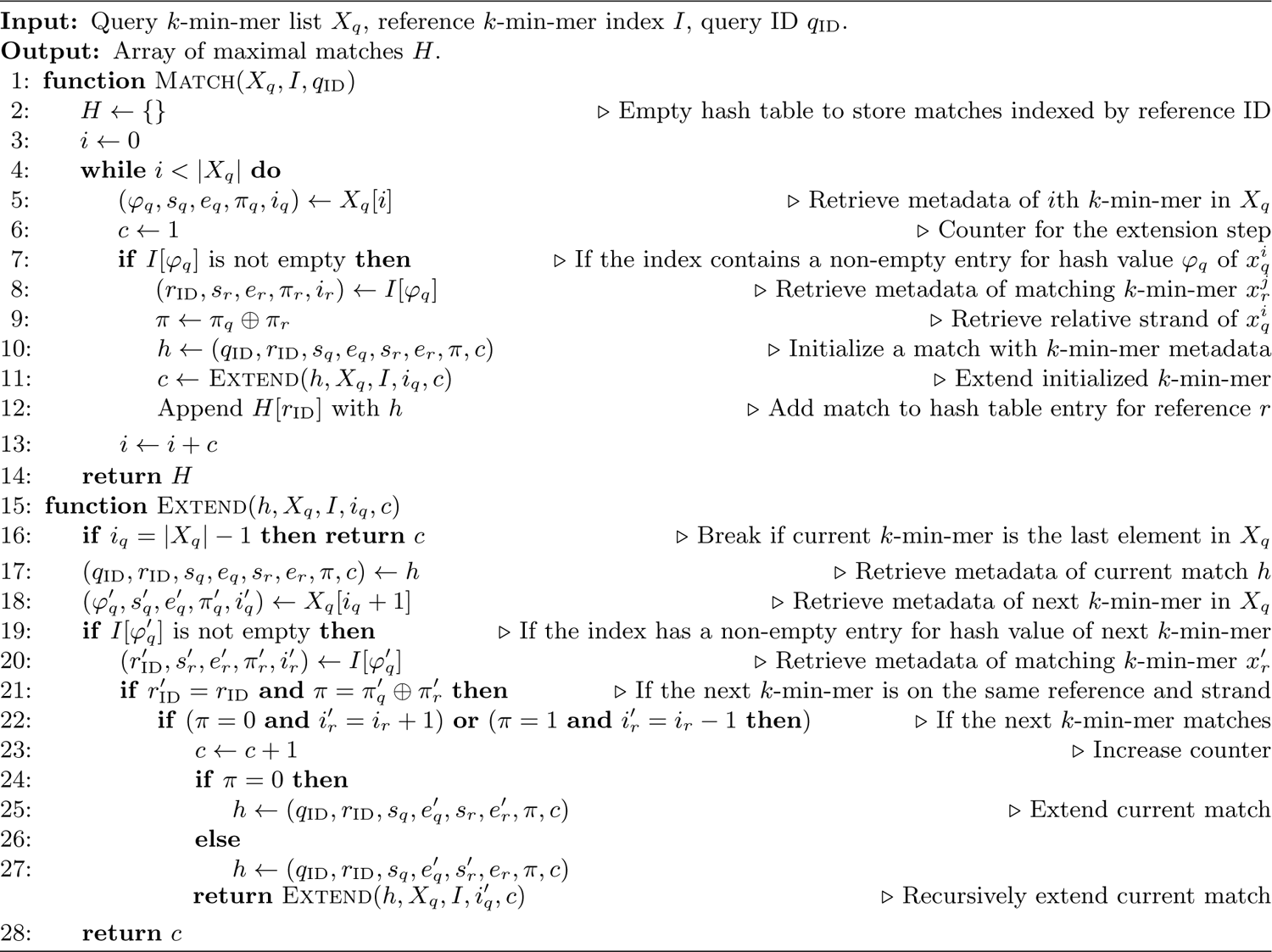

#### Algorithm 3 Pseudo-chaining *k*-min-mer matches

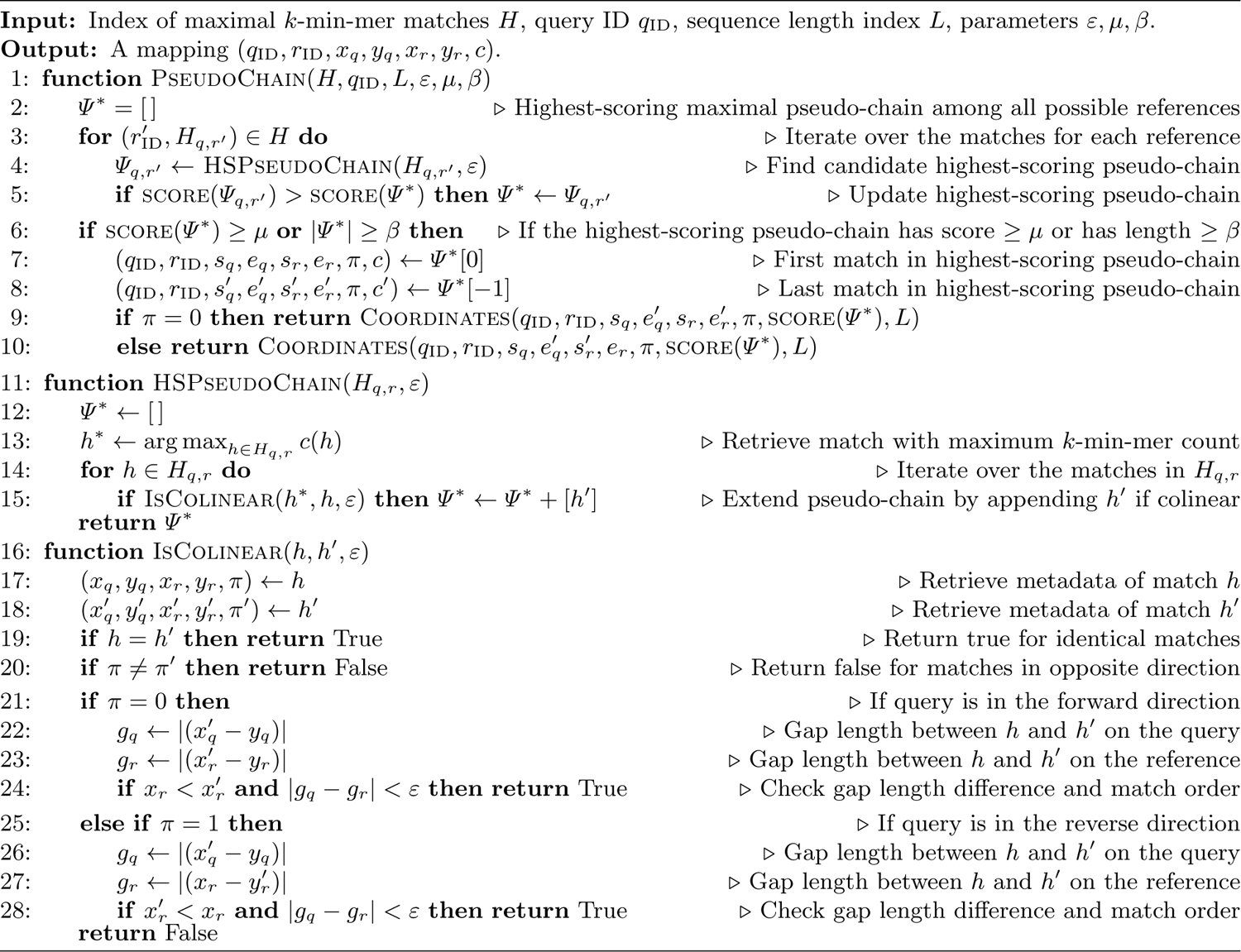

#### Algorithm 4 Obtaining final mapping coordinates

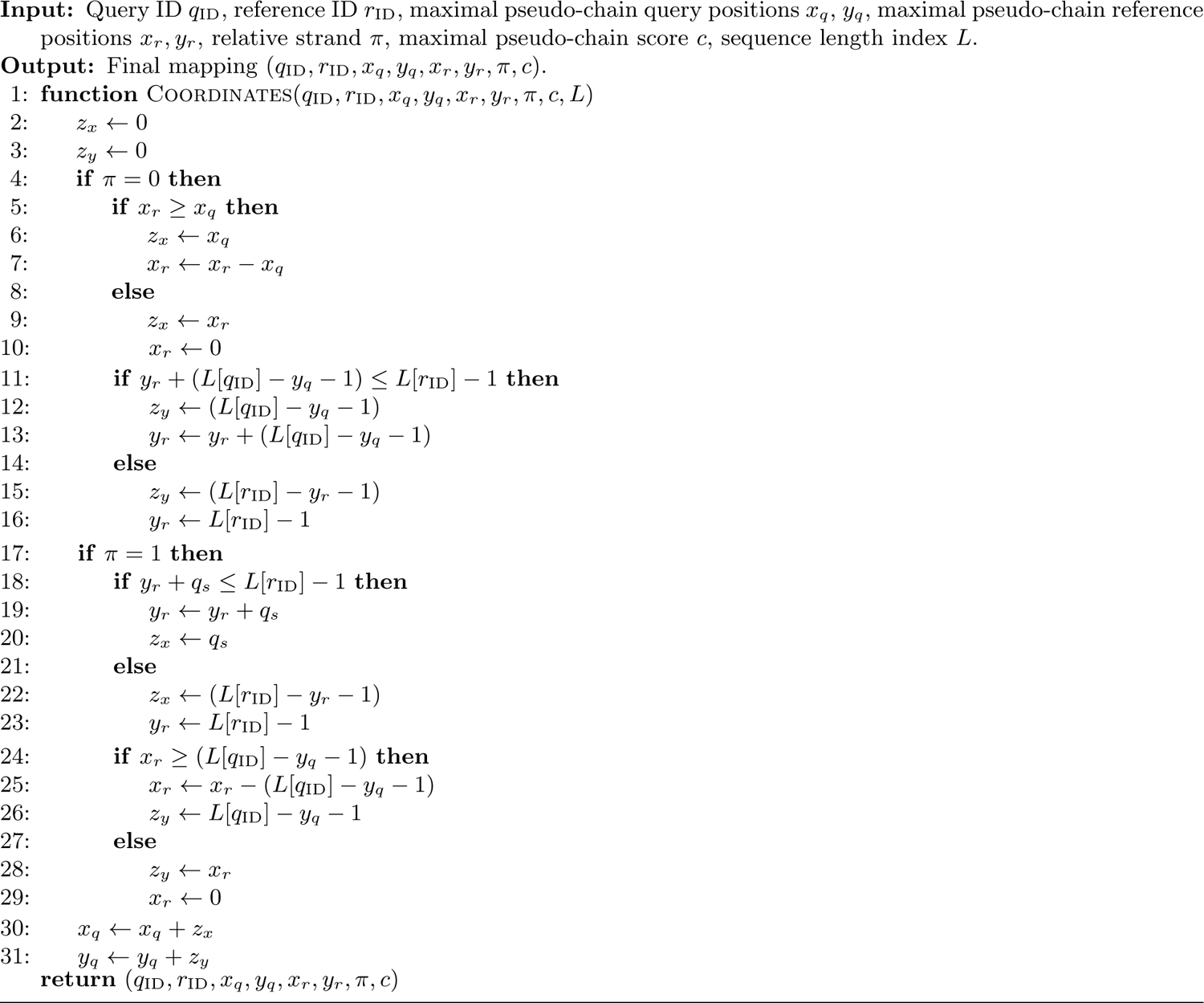

### A.2 Command lines and versions

Simulated human HiFi reads were simulated using `pbsim` at 10× coverage depth with the following command line:

~~~
ref=chm13v2.0.fa
reads=simulated-chm13v2.0-10X
pbsim $ref --model_qc PBSIM-PacBio-Simulator/data/model_qc_clr --accuracy-mean 0.99 \
       --accuracy-sd 0 --depth 10 --prefix $reads --length-mean 24000
paftools.js pbsim2fq $ref.fai “$reads”_*.maf > $reads.fa
~~~

Mappers were run using the following command lines:

~~~
mapquik $reads --reference chm13v2.0.oneline.fa --threads 10
minimap2 -x map-hifi -t 10 chm13v2.0.fa $reads
mm2-fast -x map-hifi -t 10 chm13v2.0.fa $reads
blend -t 10 -x map-hifi chm13v2.0.fa $reads
winnowmap -t 10 -W repetitive_k15.txt -x map-pb chm13v2.0.fa $reads
~~~

Tool versions:

~~~
minimap 2.24-r1122
mm2-fast 2.24-r1122
BLEND 1.0
winnowmap 2.03
mapquik 0.1.0.
~~~

### A.3 Complete simulated mapping evaluations

In this section, we report the complete results of `paftools_mapeval` evaluations on the simulated 10× coverage human genome experiment from Table 1, including the reads that were mapped at mapQ below 60. Column 2 is the mapping quality, column 3 is the number of mapped reads at this quality, and column 4 is the number of erroneously mapped reads at this quality. Column 5 is the cumulative fraction of erroneously mapped reads, and column 6 is the cumulative number of mapped reads.

`mapquik`:

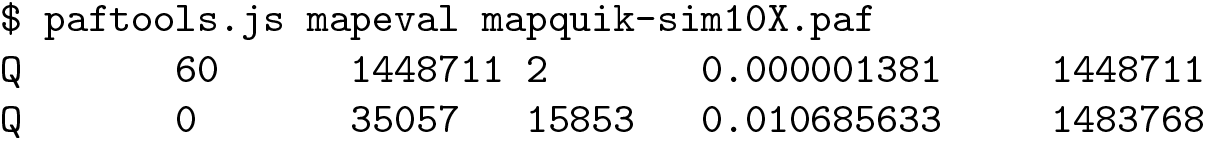

`Winnowmap2`:

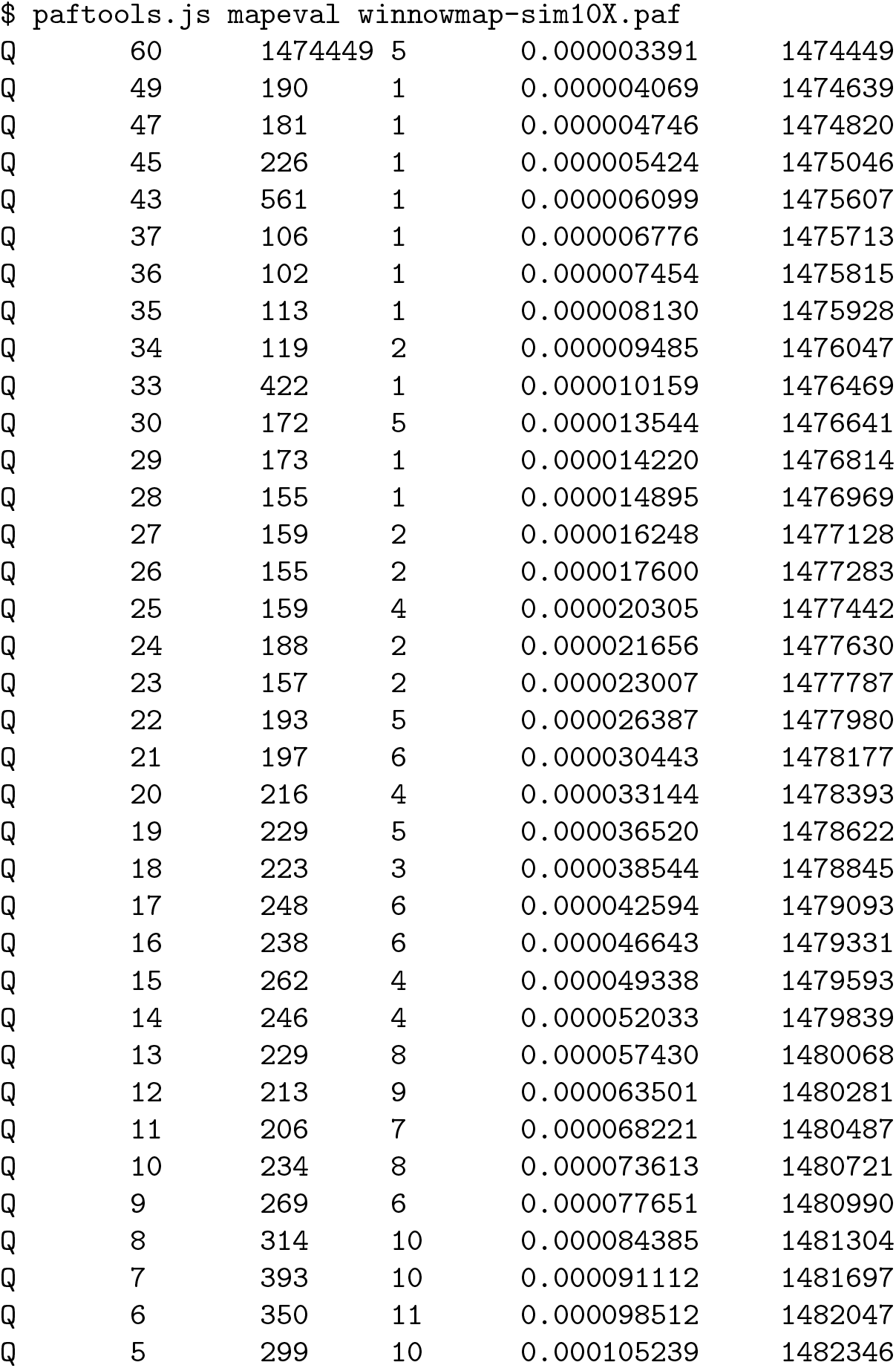
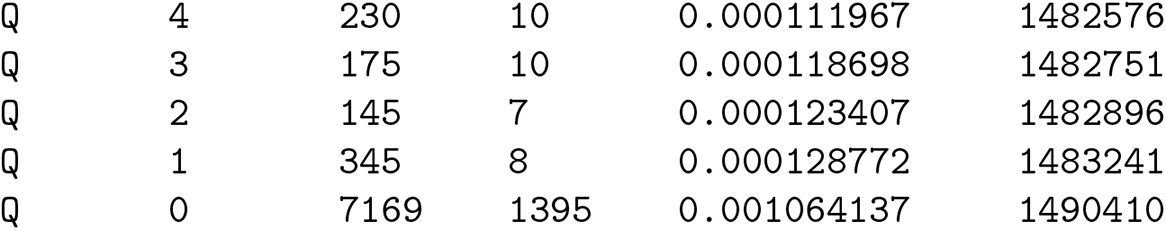

`minimap2`:

|  |  |  |  |  |  |
| --- | --- | --- | --- | --- | --- |
| Q | 60 | 1475985 | 0 | 0.000000000 | 1475985 |
| Q | 0 | 14425 | 1546 | 0.001037298 | 1490410 |

`mm2-fast`:

|  |  |  |  |  |  |
| --- | --- | --- | --- | --- | --- |
| Q | 60 | 1475985 | 0 | 0.000000000 | 1475985 |
| Q | 0 | 14425 | 1546 | 0.001037298 | 1490410 |

`BLEND`:

|  |  |  |  |  |  |
| --- | --- | --- | --- | --- | --- |
| Q | 60 | 1452658 | 1 | 0.000000688 | 1452658 |
| Q | 48 | 2033 | 1 | 0.000001375 | 1454691 |
| Q | 44 | 2050 | 1 | 0.000002059 | 1456741 |
| Q | 40 | 980 | 1 | 0.000002744 | 1457721 |
| Q | 38 | 490 | 2 | 0.000004115 | 1458211 |
| Q | 37 | 525 | 1 | 0.000004799 | 1458736 |
| Q | 36 | 516 | 1 | 0.000005482 | 1459252 |
| Q | 35 | 1011 | 1 | 0.000006163 | 1460263 |
| Q | 33 | 481 | 3 | 0.000008215 | 1460744 |
| Q | 32 | 529 | 3 | 0.000010265 | 1461273 |
| Q | 31 | 477 | 3 | 0.000012314 | 1461750 |
| Q | 30 | 495 | 2 | 0.000013678 | 1462245 |
| Q | 29 | 471 | 1 | 0.000014357 | 1462716 |
| Q | 28 | 469 | 2 | 0.000015719 | 1463185 |
| Q | 27 | 509 | 9 | 0.000021862 | 1463694 |
| Q | 26 | 469 | 9 | 0.000028002 | 1464163 |
| Q | 25 | 470 | 13 | 0.000036869 | 1464633 |
| Q | 24 | 504 | 5 | 0.000040269 | 1465137 |
| Q | 23 | 489 | 6 | 0.000044350 | 1465626 |
| Q | 22 | 527 | 7 | 0.000049108 | 1466153 |
| Q | 21 | 498 | 7 | 0.000053864 | 1466651 |
| Q | 20 | 450 | 9 | 0.000059982 | 1467101 |
| Q | 19 | 429 | 16 | 0.000070867 | 1467530 |
| Q | 18 | 478 | 15 | 0.000081062 | 1468008 |
| Q | 17 | 446 | 13 | 0.000089890 | 1468454 |
| Q | 16 | 489 | 17 | 0.000101433 | 1468943 |
| Q | 15 | 481 | 18 | 0.000113650 | 1469424 |
| Q | 14 | 511 | 18 | 0.000125856 | 1469935 |
| Q | 13 | 470 | 16 | 0.000136697 | 1470405 |
| Q | 12 | 527 | 27 | 0.000155004 | 1470932 |
| Q | 11 | 479 | 19 | 0.000167866 | 1471411 |
| Q | 10 | 576 | 33 | 0.000190219 | 1471987 |
| Q | 9 | 637 | 36 | 0.000214583 | 1472624 |
| Q | 8 | 607 | 43 | 0.000243682 | 1473231 |
| Q | 7 | 735 | 58 | 0.000282910 | 1473966 |
| Q | 6 | 715 | 52 | 0.000318035 | 1474681 |
| Q | 5 | 795 | 60 | 0.000358528 | 1475476 |
| Q | 4 | 922 | 70 | 0.000405717 | 1476398 |
| Q | 3 | 994 | 75 | 0.000456209 | 1477392 |
| Q | 2 | 1201 | 102 | 0.000524823 | 1478593 |
| Q | 1 | 1328 | 120 | 0.000605438 | 1479921 |
| Q | 0 | 10489 | 3185 | 0.002738173 | 1490410 |

### A.4 Analysis of divergence

Figure 4 shows the total number of mapped reads, the number of mapped reads at mapping quality Q60, and the number of wrongly mapped reads at Q60 of `mapquik`, when mapping simulated CHM13v2.0 reads at 10× coverage with varying levels of sequencing errors (from perfect to 10% error rate) to the CHM13v2.0 reference. For all levels, `mapquik` was run with *k* = 7, *l* = 31, *δ* = 0.01. Lowering the value of *k* for higher divergences increases sensitivity (data not shown), hence this analysis provides a pessimistic estimation of `mapquik` robustness.

**Fig. 4:**
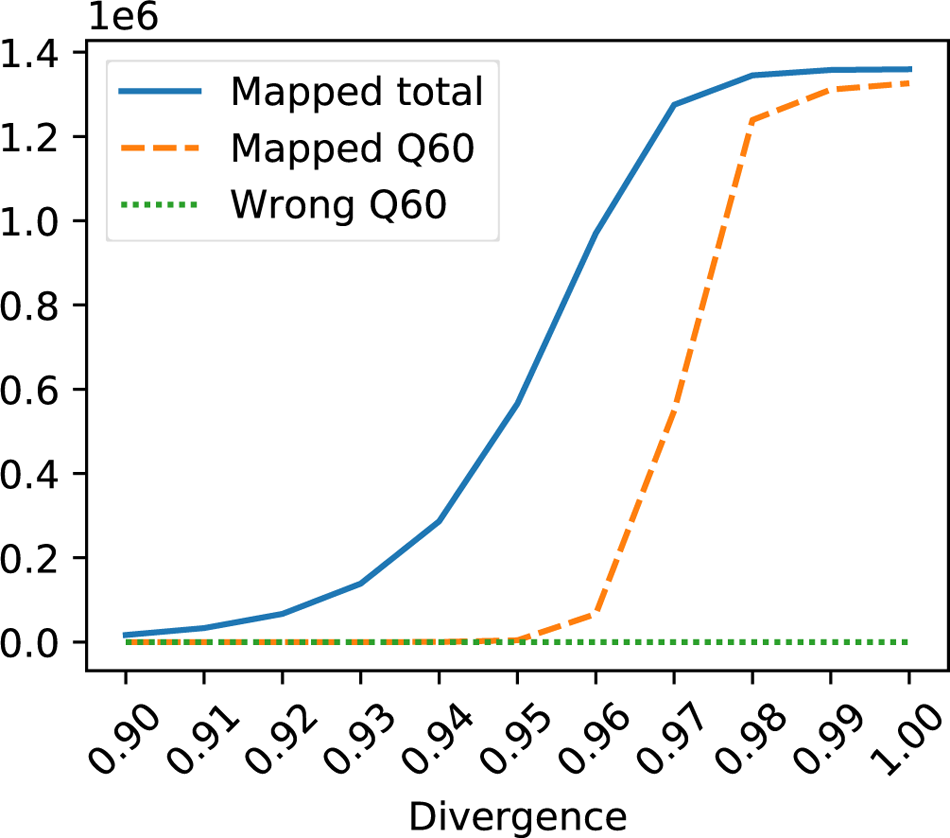
Degradation of performance of mapquik with higher divergences between reads and reference. The “Mapped total” line corresponds to the total number of mapped reads by `mapquik`. The “Mapped Q60” line corresponds to the number of reads mapped at mapping quality 60. The “Wrong Q60” line corresponds to wrongly mapped reads as assessed by `paftools mapeval`.

### A.5 Hash functions and minimizers implementation details

We use the `ntHash1` [29] method for hashing *ℓ*-mers, as implemented in https://github.com/luizirber/nthash. Minimizers are computed as universe minimizers [9], in implicit homopolymer-compressed space, i.e., all sequences are homopolymer-compressed (HPC) during minimizer computation, but positions of the HPC minimizers are reported in the original unmodified sequence.

Previous works have examined SIMD acceleration of (1) `ntHash2` [19] using AVX2 and AVX512 SIMD instructions (see `ntHash` GitHub repository pull request #9) and (2) windowed minimizer computation [43], where speed-ups of 3 − 5× were reported. The `mapquik` codebase implements several flavors of `ntHash` that can be of independent interest: A scalar (`ntHash1`) using 32- or 64-bit hashes, as well as an AVX512 implementation of `ntHash2` using 32-bit hashes. The latter ended up not being used in our tests due to the lack of an efficient compatibility with homopolymer-compressed input.

### A.6 Additional remarks on Figure 2

In the left panel of Figure 2, note that for all values of *k* shown, the distribution resembles that of a *k*-mer histogram: It follows a pseudo-normal distribution, except for the smaller peak on the left side of the curve that corresponds to erroneous pseudo-chains, which occur either due to sequencing errors in the reads, or due to hash collisions. Even though this peak is considerably smaller than that in a *k*-mer histogram (due to sampling minimizers, which may or may not have errors), we still observe a similar effect as the peak that corresponds to erroneous *k*-mers in a *k*-mer histogram: Similar to filtering out erroneous *k*-mers by imposing a threshold on the frequency of a *k*-mer (only selecting “solid” *k*-mers, for example, which have a count of *>* 1), we can filter out erroneous pseudo-chains by imposing a threshold on the score of each pseudo-chain.

